# Oxidative Modification of LHC II Associated with Photosystem II and PS I-LHC I-LHC II Membranes

**DOI:** 10.1101/2021.11.29.470042

**Authors:** Ravindra S. Kale, Jacob Seep, Larry Sallans, Laurie K. Frankel, Terry M. Bricker

**Affiliations:** Department of Biological Sciences, Biochemistry and Molecular Biology Section, Louisiana State University, Baton Rouge, LA 70803; The Rieveschl Laboratories for Mass Spectrometry, Department of Chemistry, University of Cincinnati, Cincinnati, OH 45221

**Keywords:** Photosystem I, Photosystem II, LHC II, Lhcb1, Lhcb2, tandem mass spectrometry, Reactive Oxygen Species (ROS), spinach, ^1^O_2_

## Abstract

Under aerobic conditions the production of Reactive Oxygen Species (ROS) by electron transport chains is unavoidable, and occurs in both autotrophic and heterotrophic organisms. In photosynthetic organisms both Photosystem II (PS II) and Photosystem I (PS I), in addition to the cytochrome *b*_*6*_*/f* complex, are demonstrated sources of ROS. All of these membrane protein complexes exhibit oxidative damage when isolated from field-grown plant material. An additional possible source of ROS in PS I and PS II is the distal, chlorophyll-containing light-harvesting array LHC II, which is present in both photosystems. These serve as possible sources of ^1^O_2_ produced by the interaction of ^3^O_2_ with ^3^chl* produced by intersystem crossing. We have hypothesized that amino acid residues close to the sites of ROS generation will be more susceptible to oxidative modification than distant residues. In this study, we have identified oxidized amino acid residues in a subset of the spinach LHC II proteins (Lhcb1 and Lhcb2) that were associated with either PS II membranes (i.e. BBYs) or PS I-LHC I-LHC II membranes, both of which were isolated from field-grown spinach. We identified oxidatively modified residues by high-resolution tandem mass spectrometry. Interestingly, two different patterns of oxidative modification were evident for the Lhcb1 and Lhcb2 proteins from these different sources. In the LHC II associated with PS II membranes, oxidized residues were identified to be located on the stromal surface of Lhcb1 and, to a much lesser extent, Lhcb2. Relatively few oxidized residues were identified as buried in the hydrophobic core of these proteins. The LHC II associated with PS I-LHC I-LHC II membranes, however, exhibited fewer surface-oxidized residues but, rather a large number of oxidative modifications buried in the hydrophobic core regions of both Lhcb1 and Lhcb2, adjacent to the chlorophyll prosthetic groups. These results appear to indicate that ROS, specifically ^1^O_2_, can modify the Lhcb proteins associated with both photosystems and that the LHC II associated with PS II membranes represent a different population from the LHC II associated with PS I-LHC I-LHC II membranes.

## Introduction

In higher plants (and green algae), LHC II serves as a distal antenna for both Photosystem II (PS II) and Photosystem I (PS I) (Chukhutsina et al. 2020; Croce 2020; Wientjes et al. 2013b). The chl and carotenoid prosthetic groups of these proteins absorb photons and transfer excitation energy to more proximal antennae associated with each reaction center. In this role they respond to changes in environmental illumination conditions, modulating the optical cross-sections of both photosystems. Two general types of LHC II proteins are present, those which form trimers (Lhcb1-3) and those which are monomeric (Lhcb4-6). In PS II, three different types of LHC II trimers have been designated, S(strong), M(medium) and L(loose), based on their resistance to detergent removal from the PS II core dimers (C2) (Ballottari et al. 2012). These form, with the addition of the monomeric Lhcbs, the C2S2M2 supercomplex. The monomeric Lhcbs proteins appear to transduce excitation energy from the LHC II trimers to the proximal antennae of the photosystem that is predominantly associated with the PsbB (CP47) and PsbC (CP43) proteins (Bricker 1990; Bricker and Frankel 2002). This supercomplex appears to associate with an undetermined number of L trimers (Boekema et al. 1999; Nosek et al. 2017). LHC II trimers are quite heterogeneous, containing mixtures of the Lhcb1-3 subunits. About 67% of the subunits are Lhcb1, 25% Lhcb2, and 8% are Lhcb3 (Jansson 1994).

During State 1-State 2 transitions a subpopulation of LHC II associated with PS II becomes phosphorylated, detaches from PS II, and subsequently associates with PS I. Conversely, during State 2-State 1 transitions, these components are dephosphorylated, detach from PS I and re-associate with PS II. This dynamic system serves to balance the excitation energy partitioned between the two photosystems allowing for optimal proton gradient formation and electron transport to NADP^+^. The phosphorylation and dephosphorylation of Lhcb2 appears to be particularly important with respect to state transitions (Pietrzykowska et al. 2014; Crepin and Caffarri 2015; Longoni et al. 2015). Interestingly, phosphorylation of the Lhcb1 and Lhcb2 proteins which are associated with the C2S2M2 supercomplex does not lead directly, in large measure, to the release of the S and M trimers from the core of PS II (Wientjes et al. 2013a; Crepin and Caffarri 2015; Galka et al. 2012). This suggests that the phosphorylation of these Lhcb proteins, possibly specifically associated with L trimers, may be instrumental for the association of LHC II with PS I. It should be noted that even under State 1 conditions, at least one LHC II trimer is constituently associated with the PS I-LHC I supercomplex and that, consequently, LHC II should also be considered as a component of the distal light-harvesting antenna of both PS I and PS II (Chukhutsina et al. 2020; Croce 2020; Wientjes et al. 2013b).

In higher plant PS I-LHC I-LHC II membranes (Bell et al. 2015; Bos et al. 2017) and PS I-LHC II megacomplexes (Schwarz et al. 2018), multiple LHC II trimers (up to five) can functionally associate with the PS I-LHC I supercomplex. At least one LHC II trimer appears to associate with the PsaL/PsaO region of PS I and may deliver excitation energy directly to the proximal PS I antenna which is principally associated with the PsaA and PsaB but includes PsaO and PsaL proteins (Pan et al. 2018). A second LHC II trimer may associate with the distal Lhca antenna (Benson et al. 2015). The location(s) of any additional LHC II trimers associated with the PS I-LHC I supercomplex has not yet been determined. It should be noted that in *Chlamydomonas* a subset of the monomeric LHC II proteins also appears to associate with PS I (Kargul et al. 2005; Takahashi et al. 2006; Drop et al. 2014); it is unclear if this occurs in higher plants. No monomeric Lhcb proteins were identified in the maize PS I-LHC I-LHC II structure (Pan et al. 2018), however these may have been lost during isolation of the supercomplex since one study has indicated that the monomeric Lhcb4 is associated with the maize PS I-LHC I-LHC II (Urban et al. 2020).

Crystal structures are available for LHC II trimers (*Pisum satvium*, 2.5 Å (Standfuss et al. 2005)) and cryo-EM structures are available for the LHC II proteins (trimeric and monomeric) associated with both PS II (*Arabidopsis thaliana*, 5.3 Å (van Bezouwen et al. 2017); *Spinacia oleracea*, 3.2 Å (Wei et al. 2016); *Pisum satvium*, 2.7 Å, (Su et al. 2017)) and the PS I-LHC I-LHC II supercomplex (*Zea mays*, 3.3 Å (Pan et al. 2018)).

PS II membranes have been instrumental in expanding our understanding of the structure and function of PS II. These are typically isolated by treatment of stacked thylakoid membranes with TX-100 followed by differential centrifugation (Berthold et al. 1981). These membranes exhibit high rates of oxygen evolution (generally, 400-600 μmoles O_2_•mg chl•hr), are highly enriched in PS II membrane proteins and LHC II, being almost fully devoid of PS I, the cytochrome *b*_*6*_*f* complex, and CF_1_-CF_o_. Electron microscopy indicates that individual membranes are discoidal, with a diameter of 600-1000 nm (Fig. 1b) (Møller et al. 1984; Haferkamp et al. 2010), approximating the dimensions of individual grana membranes (Austin and Staehelin 2011; Staehelin and Paolillo 2020). PS II membranes isolated in this manner are also somewhat depleted of membrane lipids (Haferkamp et al. 2010; Duchene and Siegenthaler 2000), yielding high protein packing densities (80% protein area *vs*. 70% protein packing in native grana membranes). Recently, PS I-LHC I-LHC II membranes have been isolated by treatment of stacked thylakoid membranes with a styrene-maleic acid (2:1) copolymer and differential centrifugation (Bell et al. 2015). These membranes exhibit high rates of electron transport using 2,6 dichlorophenolindophenol as an electron donor and methyl viologen as an electron acceptor (−144 μmoles O_2_•mg chl•hr), are highly enriched in PS I membrane proteins and LHC II and are almost fully devoid of other PS II components, the cytochrome *b*_*6*_*f* complex, and CF_1_-CF_o_ (Bell et al. 2015). Electron microscopy indicates that individual PS I-LHC-LHC II membranes are discoidal, with a diameter of about 113 nm ± 20 (see Fig. 1a). It has been suggested that these membranes originate from the grana margins and/or stroma lamellae (Chukhutsina et al. 2020).

**Figure 1.**
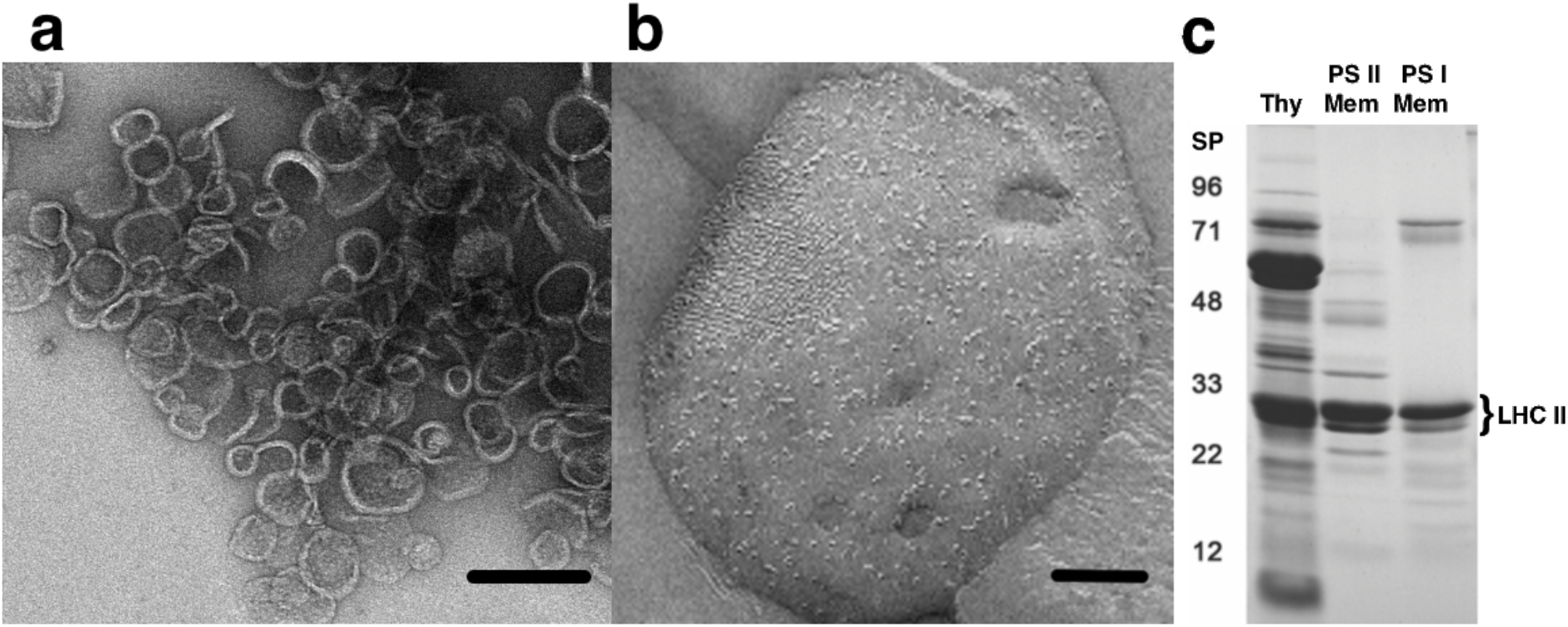
Characterization of PS II Membranes and PS I-LHC I-LHC II Membranes. a, Negatively stained electron microscopy of PS I-LHCI-LHC II membranes. These present as membrane disks with a diameter of 113 ± 20 nm (n=22), scale bar = 200 nm. b, Freeze-fracture, negatively stained image of PS II membranes with a diameter of about 800 nm, scale bar = 200 nm. This panel was adapted from Fig. 1D of (Haferkamp et al. 2010). c, LiDS-PAGE of Thylakoids, PS II membranes, and PSI-LHCI-LHCII membranes. Individual lanes were loaded with 4 µg chl. The PSI-LHCI-LHCII membranes are highly enriched in PSI core proteins, contain significant amounts of LHC II (indicated to the right), and are highly depleted of PS II components (BBY) as previously reported (Bell et al. 2015). THY, thylakoids; PS II Mem, PS II membranes (BBYs), PSI Mem, PSI-LHCI-LHCII membranes;. Apparent molecular masses of the standard proteins (SP) are shown to the left. This figure was redrawn from (Bell et al. 2015).

Under aerobic conditions the production of Reactive Oxygen Species (ROS) by electron transport chains is unavoidable, occurring in both autotrophic and heterotrophic organisms (Foyer 2018; Munro and Treberg 2017). All of the photosynthetic membrane protein complexes (PS II, the *b*_*6*_*/f* complex, and PS I) can produce ROS under specific conditions. An additional possible source of ROS in PS II and PS I are the chlorophyll-containing, light-harvesting arrays present in both photosystems. These may produce ^1^O_2_ by the interaction of ground-state ^3^O_2_ with the ^3^chl* via intersystem crossing. The production of ^1^O_2_ is potentially quite harmful as this ROS can oxidatively damage proteins, pigments, lipids and nucleic acids (Halliwell 2006). Earlier, we had shown that LHC I (Lhca1-4), the distal light-harvesting array associated with PS I, exhibited extensive oxidative modification (Kale et al. 2020) which was consistent with the production of ROS (probably ^1^O_2_) by LHC I. Interestingly, very few oxidative modifications were observed associated with the extensive proximal antenna of PS I associated with PsaA and PsaB subunits.

In the current study, high-resolution tandem mass spectrometry was used to identify the location of oxidatively modified residues within the LHC II populations associated with both PS II membranes and PS I-LHC I-LHC II membranes. These were both isolated from field-grown spinach. The identified modified residues were mapped onto the Lhcb1 and Lhcb2 subunits of the LHC II trimer associated with the 3.3 Å resolution *Zea mays* PS I - LHC I - LHC II structure (Pan et al. 2018). Previously, we had used these methods to identify natively oxidized residues in spinach PS II (Frankel et al. 2012, 2013b; Frankel et al. 2013a) (and PS II residues modified during photoinhibition (Kale et al. 2017; Kumar et al. 2021)), the cytochrome *b*_*6*_*/f* complex (Taylor et al. 2018) and the PSI-LHC I supercomplex (Kale et al. 2020).

## Materials and Methods

Photosystem II membranes and PS I-LHC I-LHC II-enriched membranes were prepared as previously described (Berthold et al. 1981; Bell et al. 2015). LiDS-PAGE (Delepelaire and Chua 1979) was performed using a non-oxidizing gel system (Rabilloud et al. 1995). Electrophoresis was stopped when the proteins entered the resolving gel ∼1.0 cm, the gel was stained with Coomassie blue, destained, and the large protein bands containing the LHC II proteins associated with either PS II membranes or PS I-LHC I-LHC II membranes were harvested. Standard “in gel” digestion protocols were used for both trypsin and chymotrypsin proteolysis. These experiments were performed in duplicate.

For mass spectrometry, the proteolytic peptides were isolated from the gel fragments and resolved by high performance liquid chromatography on a C:18 reversed-phase column. Chromatography conditions were as described in (Frankel et al. 2012); data were collected during the acetonitrile gradient and during the wash phase (80% acetonitrile) of the C:18 column. Electrospray ionization was used to introduce the resolved proteolytic peptides into a Thermo Scientific Orbitrap Fusion Lumos mass spectrometer. The samples were analyzed in a data-dependent mode with one Orbitrap MS^1^ scan acquired simultaneously with up to ten linear ion trap MS^2^ scans. The Thermofisher RAW files were converted to MGF files and analyzed by the MassMatrix Program (Xu and Freitas 2009), which identified the individual proteolytic peptides and their parent proteins, and allowed localization of oxidative modifications. A FASTA library containing the mature sequences of the nine spinach Lhcb proteins (Lhcb1.1-1.3, Lhcb2-6) was searched, as was a decoy library containing the reversed amino acid sequences of the Lhcbs. For protein identification, no positive hits to this decoy library were allowed (decoy hits = 0.00%). Twelve different types of oxidative modifications were considered during peptide identification, with a maximum of two modifications being allowed per peptide. Extremely rigorous p-value thresholds were used for preliminary identification of oxidized peptides with *pp-tag* value of ≤ 10^−5^ and either *pp* or *pp2* being ≤10^−5^ (Xu and Freitas 2009; Bricker et al. 2015; Kale et al. 2017). The MS^2^ spectra of the peptides exhibiting these *p*-value thresholds were then examined manually with the quality of the data being confirmed. Only peptides with charge states of ^+^3 or lower were considered, as many of the collision-induced dissociation fragments obtained from higher ionization state peptides were indistinguishable from noise in the MS^2^ spectra. The allowed mass error of the precursor MS^1^ ion was required to be ≤ 5.0 ppm. The identified oxidized amino acid residues meeting all of these very conservative identification conditions were mapped onto the cryo-EM structure of the LHC II trimer associated with the 3.3 Å resolution *Zea mays* PS I-LHC I-LHC II structure (Pan et al. 2018) using PYMOL (DeLano 2002). This trimer contains two copies of Lhcb1 and a single copy of Lhcb2.

For negative stain electron microscopy, samples of PSI-LHCI-LHCII membranes were used directly by negative staining with 2% uranyl acetate (UA) on a glow discharged carbon-coated copper grid. Electron microscopy was performed on a JEOL 1400 TEM equipped with an LaB_6_ cathode, and an operating voltage of 120 kV. Images were recorded at 60,000 x magnification with the Å/pixel 1.73Å, binning the images to 2048×2048 pixels using a bottom mount water-cooled CCD-camera with high resolution (Gatan, Pleasanton, CA, USA).

## Results and Discussion

Photosystem II membranes and PS I-LHC I-LHC II membranes are dramatically different in size (Fig. 1a and 1b). Fig. 1a illustrates the appearance of PS I-LHC I-LHC II membranes negatively stained with uranyl acetate. These are quite consistent in size with a diameter of 113 ± 20 nm (n=22). Interestingly, these are very similar to the size and appearance of partially styrene maleic acid-solubilized membranes from other biological sources (Angelisová et al. 2019; Zhang et al. 2017). Fig. 1b illustrates the appearance of freeze-fractured PS II membranes (this image was reproduced from (Haferkamp et al. 2010), their Fig. 1D; for a TEM image of these PS II membranes, please consult (Møller et al. 1984). These are large, 600-1000 nm in diameter, and approximate the size of individual grana lamella (Austin and Staehelin 2011; Staehelin and Paolillo 2020). The protein composition of these membranes was indistinguishable from our previous reports (Bell et al. 2015; Kale et al. 2020). The PS II membranes are highly enriched in PS II core components (PsbA, PsbB, PsbC, PsbD, etc.) and LHC II. The PS I-LHC I-LHC II membranes are highly enriched in PS I core components (PsaA, PsaB, PsaC, PsaD, etc.), Lhca1-4, and LHC II (Bell et al. 2015; Kale et al. 2020). We had earlier shown that both types of membranes were also strongly depleted of cytochrome *b*_*6*_*/f* and CF_1_-CF_o_ subunits (Bell et al. 2015). The LHC II associated with the PS I-LHC I supercomplex was functionally connected to the reaction center, increasing its optical cross section (Bell et al. 2015) and transferring excitation energy relatively slowly from LHC II to P_700_ but with a 94% efficiency (Bos et al. 2017). Interestingly, multiple LHC II trimers were present per PS I reaction center (Bell et al. 2015; Bos et al. 2017; Schwarz et al. 2018). In both PS II membranes and PS I-LHC I-LHC II membranes there was no evidence for the presence of significant populations of LHC II which were not functionally connected to the photosystems.

Spinach contains at least eight Lhcb proteins, Lhcb1.1-Lhcb1.3, Lhcb2, Lhcb3, Lhcb4-6. In this study we examined oxidative modifications in Lhcb1 and Lhcb2 proteins which are found in LHC II trimers. Lhcb3, while also present in LHC II trimers, only constitutes about 8% of the Lhcb proteins present (Jansson 1994). In our hands, this low abundance precluded the detection of oxidative modifications on Lhcb3 by mass spectrometry. When compared to the other Lhcb proteins, the monomeric Lhcb4-6 proteins also have relatively low abundance and were not examined in this study. In Fig. 2a, the sequences of these proteins are shown. Only slight sequence differences (< 4%) are found among the Lhcb1.1-Lhcb1.3 (henceforth Lhcb1) isoforms; consequently, we did not attempt to distinguish between these different molecular species. The sequence differences between Lhcb1 and Lhcb2, however, are more substantial (>15%, Fig. 2a, highlighted in cyan), and were sufficient, in the vast majority of instances, to distinguish between oxidative modifications of Lhcb1 and Lhcb2. If a particular oxidative modification was present on a peptide which was common to both Lhcb1 and Lhcb2 (observed extremely infrequently), then the peptide in question was eliminated from further analysis. Fig. 2b shows a phylogenetic tree, illustrating the relationship between the trimer-forming Lhcbs (Lhcb1-3) and the monomeric Lhcbs (Lhcb4-6). The Lhcb1 isoforms could not be discriminated from each other in this analysis and the close relationship of Lhcb2 and Lhcb3 to Lhcb1 is evident. The monomeric Lhcbs are significantly more distantly related to Lhcb1-3. For this phylogenetic tree the sequence of spinach Lhca1 was used as an outgroup.

**Figure 2.**
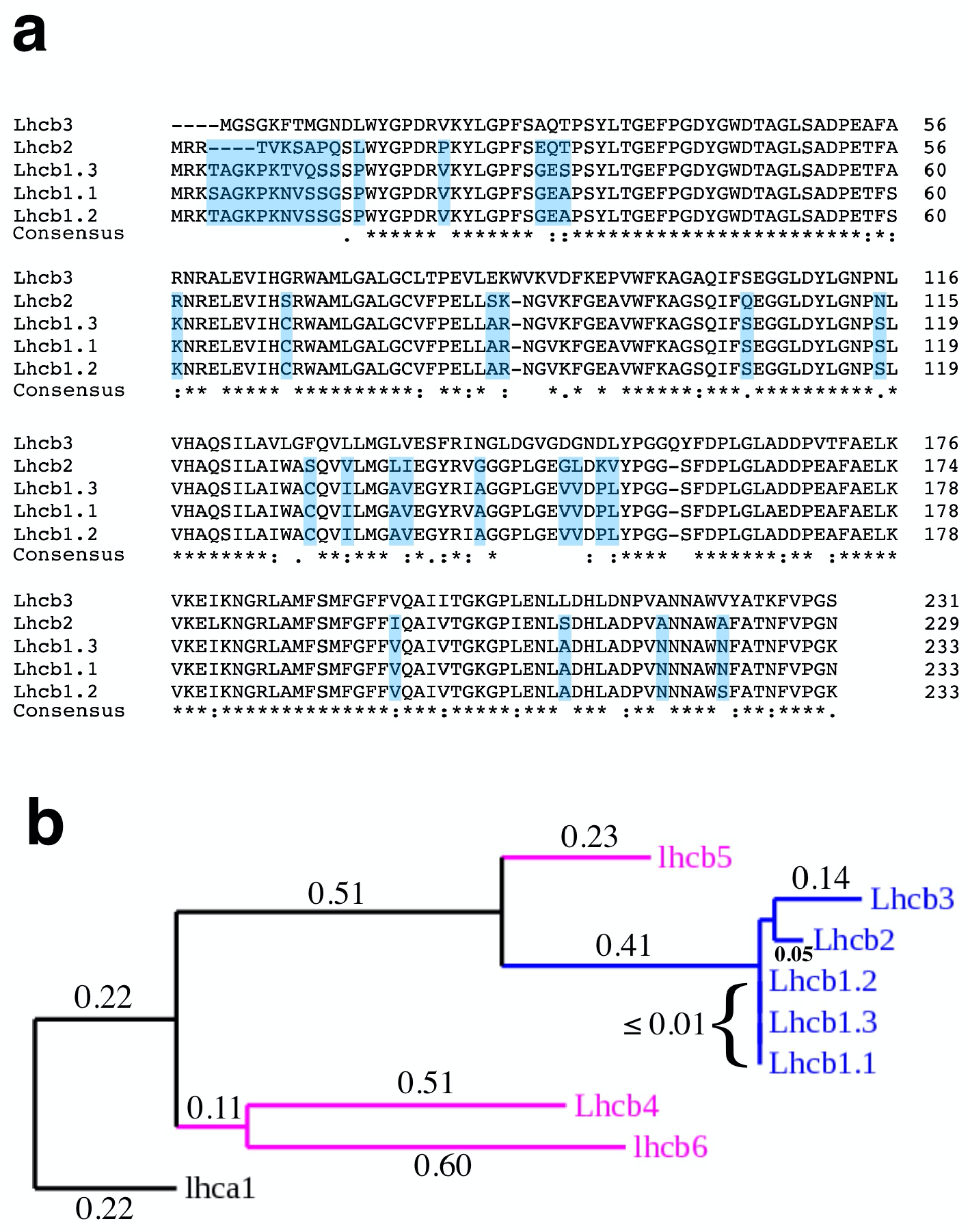
Sequence Comparison of the Trimer-Forming Lhcbs from Spinach. a, Lhcb 1.1-1.3, 2 and 3. Residues which allow differentiation between the different Lhcb1 isoforms and Lhcb2 are highlighted in cyan. Clustal Omega (Sievers and Higgins 2018) was used to align the mature sequences of these proteins. b, Phylogenetic Tree Comparing the Spinach Lhcb Proteins. For completeness, the mature sequences of the monomeric Lhcbs, Lhcb4-6, were included in this analysis. Spinach Lhca1, which is present in the distal antenna for PS I (Lhca1-Lhca4), was included as the outgroup. The trimeric Lhcbs are shown in blue, the monomeric Lhcbs are shown in magenta, and the outgroup Lhca1, is shown in black. Branch lengths are shown numerically (Dereeper et al. 2008; Castresana 2000; Anisimova and Gascuel 2006; Chevenet et al. 2006).

The Lhcb proteins are quite hydrophobic, which often poses challenges for tandem mass spectrometry experiments. Hydrophobic peptides are more difficult to elute from standard (C:18) reversed-phase chromatography matrixes and ionize more poorly than hydrophilic peptides (Dupree et al. 2020). In our experiments, high sequence coverage (∼90%) was obtained for all examined subunits of the LHC II present in both PS II membranes and PS I-LHC I-LHC II membranes (Table 1). We used trypsin and chymotrypsin, individually, for proteolytic digestion as this yielded a larger portfolio of candidate peptides for analysis. Additionally, the analysis of peptides eluted during the wash phase (80% acetonitrile) of the chromatographic elution yielded more hydrophobic peptides for analysis. Finally, oxidative modification of hydrophobic peptides decreases their hydrophobicity, decreasing their retention on reversed-phase columns and increasing their potential for elution and ionization. Over 14,000 tandem mass spectra (MS^2^) were examined for PS II membranes, and over 8000 spectra were examined for PS I-LHC I-LHC II membranes. Tandem mass spectrometry analysis of the tryptic and chymotryptic peptides of the Lhcb proteins associated with PS II membranes and PS I-LHC I-LHC II membranes indicated 35 residues were oxidatively modified in the Lhcb proteins associated with PS II membranes and that 39 residues were oxidized in the Lhcb proteins associated with the PS I-LHC I-LHC II membranes (Table 2). While the total number of oxidative modifications was similar in the LHC II isolated from both sources, the distribution of these oxidative modifications was quite different (see below).

**Table 1.**
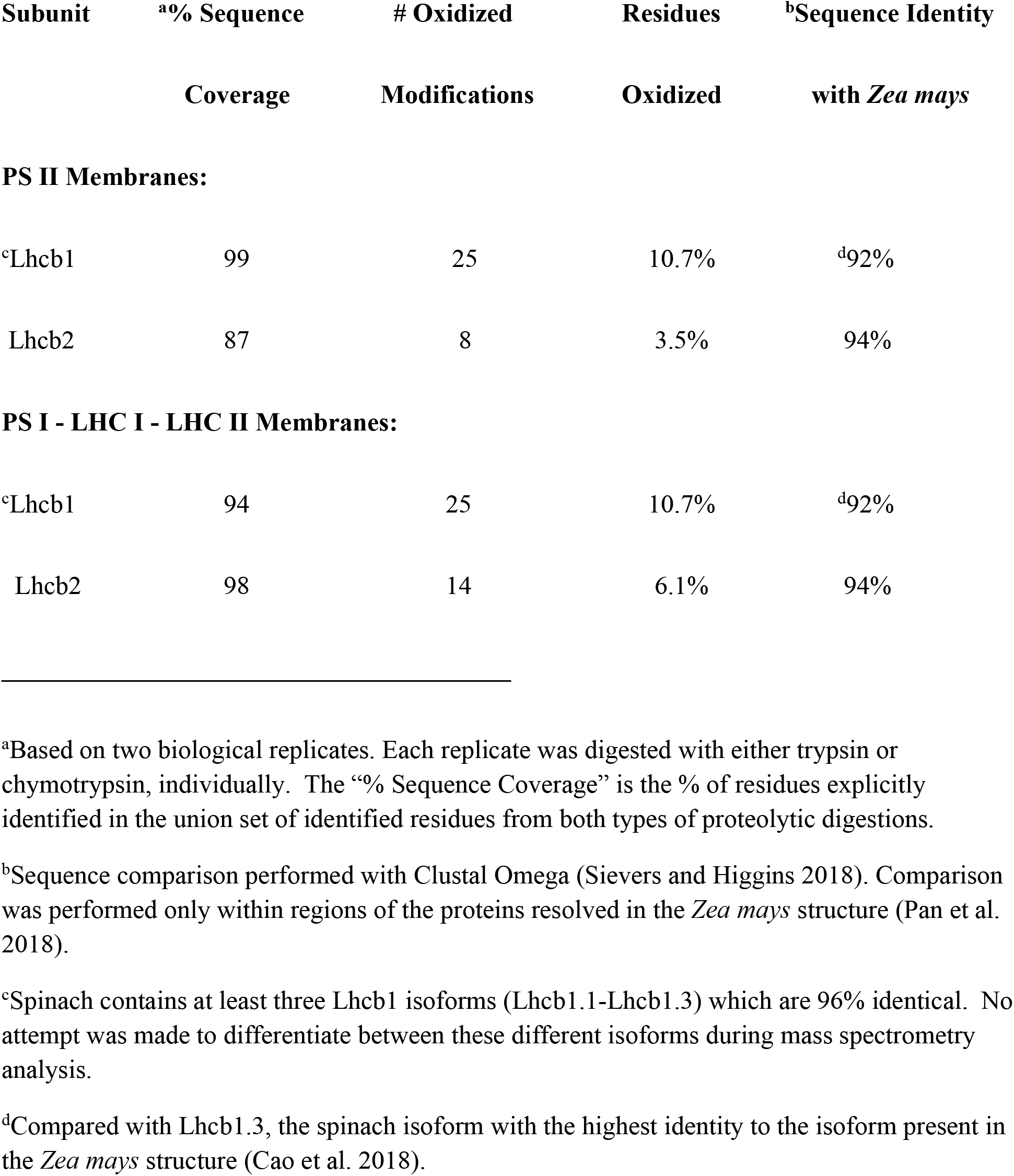
Summary Data for Mass Spectrometry and Sequence Comparison of Spinach Lhcb1 and Lhcb2 to *Zea mays* Lhcb1 and Lhcb2.

| Subunit | <sup>a</sup>% Sequence Coverage | # Oxidized Modifications | Residues Oxidized | <sup>b</sup>Sequence Identity with *Zea mays* |
| --- | --- | --- | --- | --- |
| **PS II Membranes:** |  |  |  |  |
| <sup>c</sup>Lhcb1 | 99 | 25 | 10.7% | <sup>d</sup>92% |
| Lhcb2 | 87 | 8 | 3.5% | 94% |
| **PS I - LHC I - LHC II Membranes:** |  |  |  |  |
| <sup>c</sup>Lhcb1 | 94 | 25 | 10.7% | <sup>d</sup>92% |
| Lhcb2 | 98 | 14 | 6.1% | 94% |

<sup>a</sup>Based on two biological replicates. Each replicate was digested with either trypsin or chymotrypsin, individually. The “% Sequence Coverage” is the % of residues explicitly identified in the union set of identified residues from both types of proteolytic digestions.
<sup>b</sup>Sequence comparison performed with Clustal Omega (Sievers and Higgins 2018). Comparison was performed only within regions of the proteins resolved in the *Zea mays* structure (Pan et al. 2018).
<sup>c</sup>Spinach contains at least three Lhcb1 isoforms (Lhcb1.1-Lhcb1.3) which are 96% identical. No attempt was made to differentiate between these different isoforms during mass spectrometry analysis.
<sup>d</sup>Compared with Lhcb1.3, the spinach isoform with the highest identity to the isoform present in the *Zea mays* structure (Cao et al. 2018).

**Table 2.**
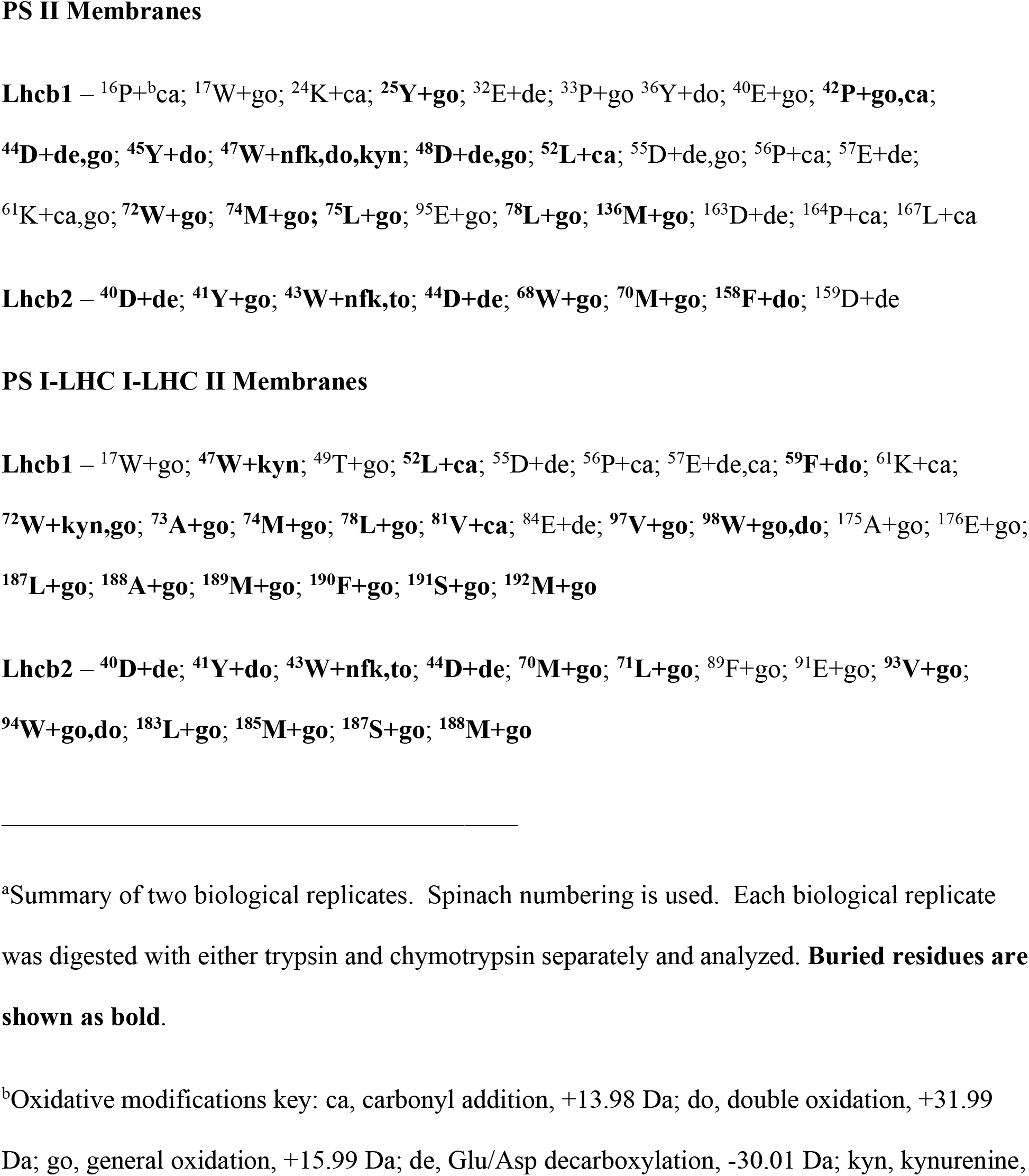
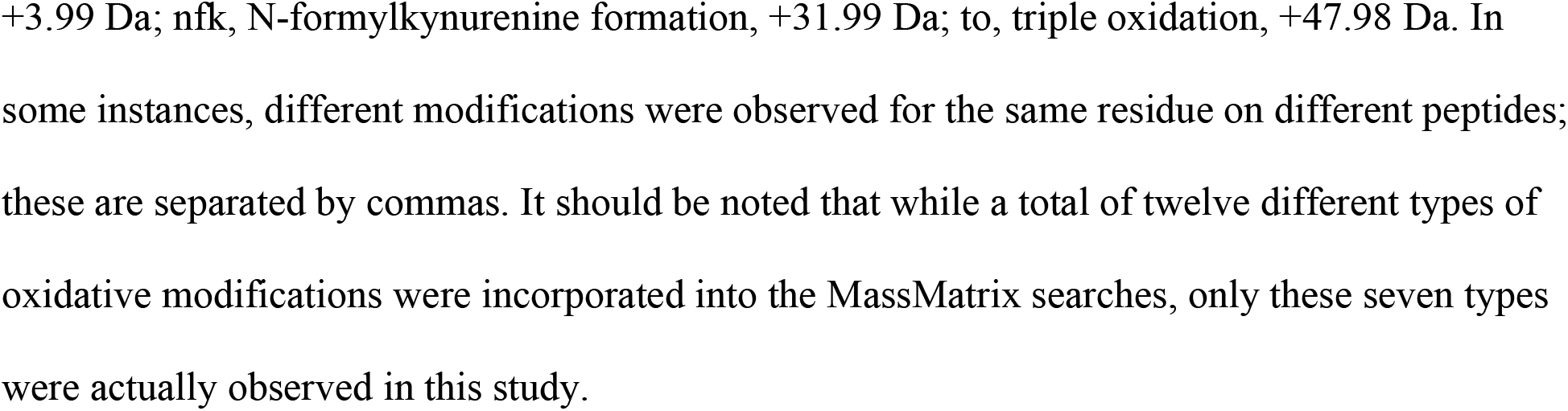
^a^Natively Oxidized Residues in Lhcb1 and Lhcb2 Isolated from PS II Membranes and PS I-LHC I-LHC II Membranes.

Fig. 3 shows the quality of the data used for the identification of oxidized amino acid residues in this study. The tandem mass spectrometry data collected for the M^+2^ tryptic peptide ^72^W-^88^R of Lhcb1 is illustrated. This peptide bears a general oxidation modification (mass change +16 Da) at ^74^M. The observed mass accuracy for the parent ion was -0.00 ppm. The *pp, pp*_*2*_ and *pp*_*tag*_ values for this peptide were 10^−3.1^, 10^−6.1^ and 10^−5.2^ (Xu and Freitas 2009), respectively, and thus was among the lowest quality peptides used in this study. Even this lower quality peptide, however, exhibited nearly complete y- and b-ion series. This allowed identification of the peptide and localization of the oxidative mass modification. This result demonstrates that the use of *p* values ≤ 10^−5^ provided high quality peptide identifications for the localization of these post-translational modifications.

**Figure 3.**
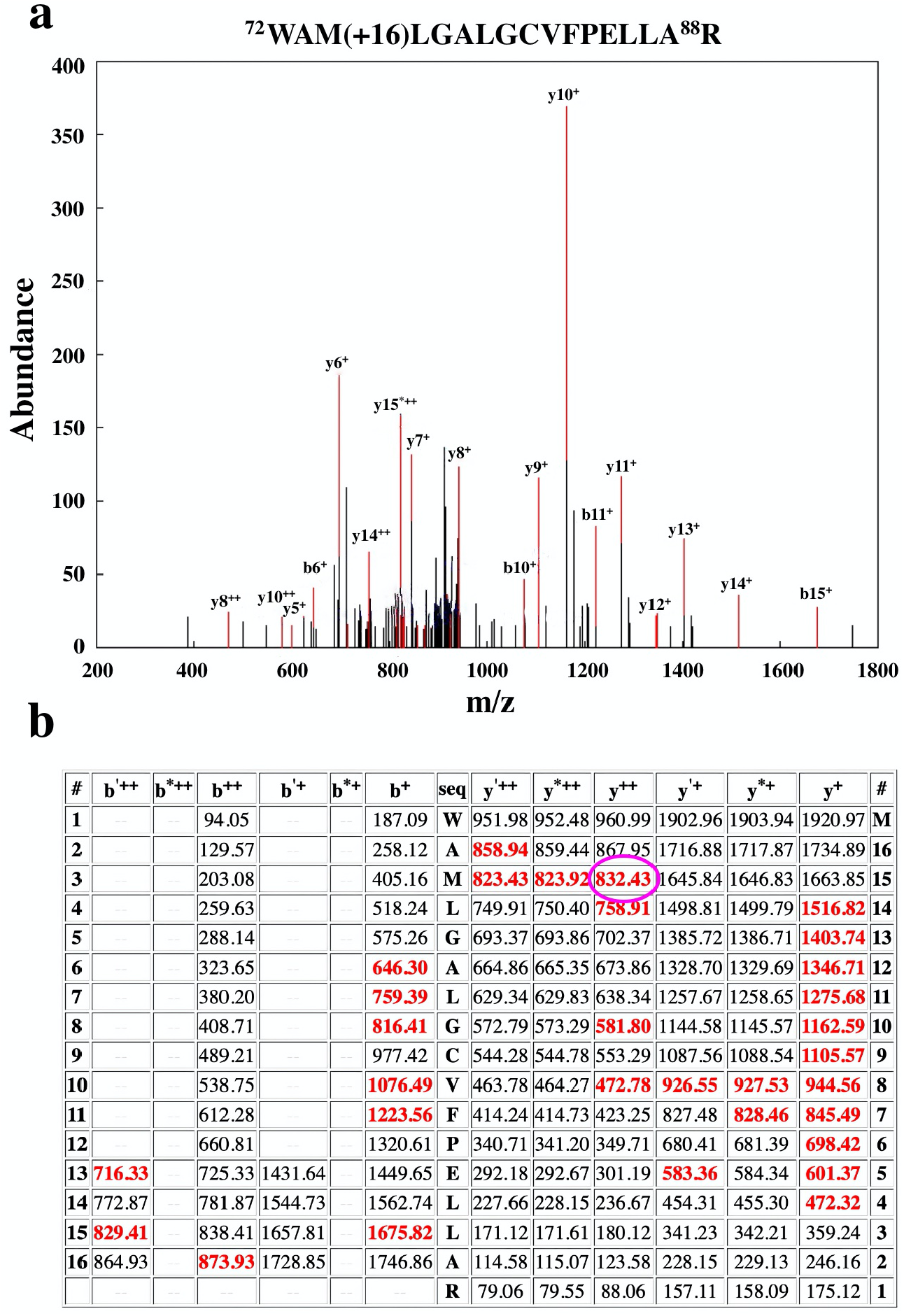
Quality of the Tandem Mass Spectrometry. Shown is the mass spectrometry result for the tryptic peptide Lhcb1:^72^W-^88^R which contains a modified ^74^M + go (+16 amu) modification. a, The CID spectrum of the modified peptide Lhcb1:^72^WAM(+16)LGALGCVFPELLA^88^R. Subsets of the identified ions are labeled. b, Table of all predicted masses for the y- and b-ions generated from this peptide sequence. Ions identified in the CID spectrum above are shown in red. The b’^++^, b’^+^ y’^++^ and y’^+^ ions are generated by the neutral loss of water while the b*^++^, b*^+^ y*^++^ and y*^+^ ions are generated from the loss of ammonia. Considering only the y- and b-ions, the ions y15^++^ (magenta ellipse), b6^+^-b8^+^ and b10^+^-b11^+^, and b16^++^ all exhibit a +16 Da mass modification. This verifies that ^74^M contains an oxidative modification which adds 16 Da to the methionyl residue. The *p*-values for this peptide were *pp* = 10^−3.1^, *pp*_*2*_ = 10^−6.1^ and *pp*_*tag*_ = 10^−5.2^ (Xu and Freitas 2009). The observed mass accuracy for the parent ion (MS^1^) was -0.00 ppm (predicted mass of peptide = 1920.9710 Da, observed mass of peptide = 1920.9710).

The identity of these oxidized residues and the types of modifications observed are presented in Table 2. It should be noted that it is very unlikely that all of the observed modifications would be present on every copy of an examined Lhcb present in a given sample. As we have noted previously (Kale et al. 2020), “*ROS modification of a particular amino acid residue is a stochastic rather than a mechanistic process. The probability that a residue will be modified depends on the ROS type and concentration, the susceptibility of a residue to a particular ROS type, the residency time of the ROS in proximity to the residue, and the lifetime of the ROS species. Our working hypothesis is that residues located near a site of ROS production (or along an ROS exit pathway) have a higher probability of oxidative modification than those distant from an ROS production site (or not along an ROS exit pathway)*.*”* Consequently, these detectable modifications are present within the **<u>populations</u>** of Lhcb1 and Lhcb2 present in our biological samples. In our study, the PS II membranes and PS I-LHC I-LHC II membranes were isolated from field-grown market spinach. The growth conditions, while unknown, would likely include the variety of abiotic stress factors typical of field grown plants (high and/or fluctuating light intensities, herbivory, high and/or low temperature variation, nutrient limitations, etc.) (Choudhary et al. 2017). These environmental factors increase ROS production. Consequently, the oxidative modifications which we observe are probably typical of those found in the field environment and are probably more extreme than those that would be observed in laboratory-grown plants cultivated under controlled environmental conditions.

In PS II, four LHC II trimers associate with each C2 core complex, forming the C2S2M2 supercomplex. Two S2 trimers are associated strongly with the core complex while two M2 trimers associate more weakly. Additionally, an indeterminate number of loosely bound trimers (Lx) associate with each C2S2M2 supercomplex. The location(s) of these Lx trimers with respect to the C2S2M2 supercomplex have not been determined. Numerous cryo-EM structures are available for the C2S2M2 supercomplexes isolated from a number of species including *Pisum satvium, Spinacia oleracea* and *Arabidopsis thaliana* (Su et al. 2017; Wei et al. 2016; van Bezouwen et al. 2017). No high-resolution structure is available for the spinach PS I-LHC I-LHC II supercomplex. However, a structure is available for the *Zea mays* supercomplex which contains a single LHC II trimer (PDB: 5ZJI,(Pan et al. 2018)). This trimer is centered on the PsaK-PsaO axis and was modeled to contain two copies of Lhcb1 and a single copy of Lhcb2. It should be noted that larger numbers of LHC II trimers are believed to associate with the PS I-LHC I supercomplex *in vivo*. PS I-LHC I-LHC II membranes contain 1-5 functionally coupled LHC II trimers depending on the particular biochemical isolate examined (Bell et al. 2015; Bos et al. 2017; Schwarz et al. 2018). In *Chlamydomonas*, cryo-EM imaging indicated that two LHC II trimers are associated with the PS I-LHC I supercomplex in State 2 (Huang et al. 2021), with one LHC II trimer (LHCII-1) being approximately positioned as observed in *Z. mays* (Pan et al. 2018) while the second trimer, LHCII-2, associates with both LHCII-1 and Lhca2 of the LHC I distal antenna. In Arabidopsis, a cryo-EM study indicated that one LHC II trimer was positioned as observed in *Z. mays* (Pan et al. 2018) and a second LHC II appeared to be associated with the Lhca2 and Lhca4 subunits (and/or Lhca2 and Lhca3) of LHC I (Yadav et al. 2017). The locations of any additional LHC II trimers which associate with the PS I-LHC I supercomplex have not been determined.

The sequence identity between spinach Lhcb1 and Lhcb2 and the *Z. mays* subunits is quite high (92-94%) (Table I). This high degree of identity allowed us to map the oxidized amino acids that we observed on the spinach proteins onto the crystal structure of the *Z. mays* LHC II trimer associated with the PS I-LHC I-LHC II supercomplex (Pan et al. 2018), which was modeled to contain two copies of Lhcb1 and one copy of Lhcb2. We used this structure as a template to map the oxidized residues which we observed for both the LHC II associated with PS II membranes and PS I-LHC I-LHC II membranes (Fig. 4). In the ^1^LHC II associated with PS II membranes, 38% of the oxidatively modified residues were surface-localized while in PS I-LHC I-LHC II membranes, 28% of the oxidized residues were surface-localized. In both instances, most of the surface-localized oxidatively modified residues are on the stromal-exposed surfaces of the Lhcb1 and Lhcb2 proteins. The identification of surface-exposed oxidatively modified residues was not unexpected. As we have noted earlier (Kale et al. 2020), the ROS produced by PS II, the cytochrome *b*_*6*_ *f* complex, and PS I may be released to the stromal (or lumenal) compartments depending on the ROS source. If this ROS escapes detoxification (Tripathy and Oelmüller 2012; Das and Roychoudhury 2014) it may contribute to the oxidative modification of the surface-exposed domains of photosynthetic membrane protein complexes, including LHC II trimers.

**Figure 4.**
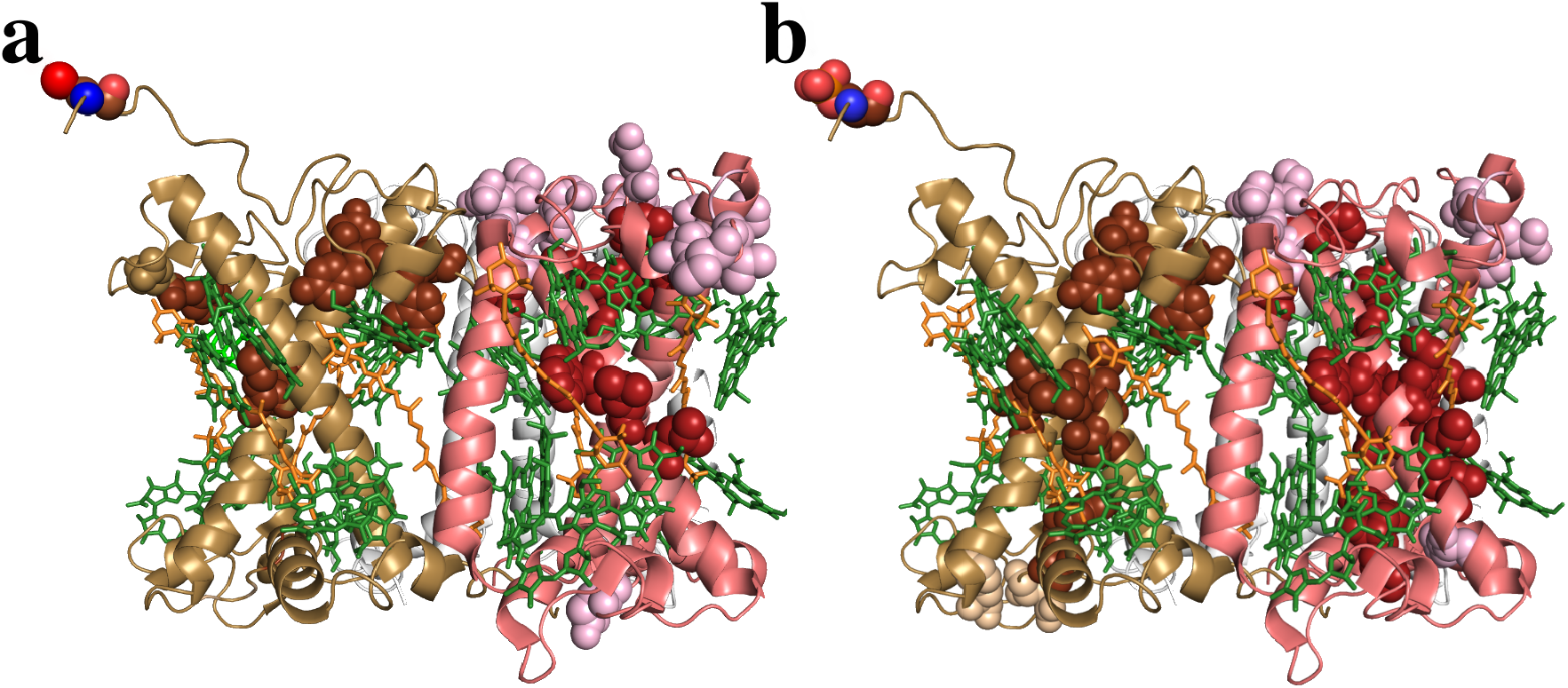
Overview of Natively Oxidized Amino Acid Residues in the Spinach LHC II associated with either a, PS II membranes, or b, PS I-LHC I-LHC II membranes. a, PS II membranes - viewpoint, within the plane of the membrane, of Lhcb2 (light brown) and Lhcb1 (pink) associated with an LHC II trimer. A second Lhcb1 monomer (white) is in the background and largely hidden. b, PS I-LHC I-LHC II membranes – viewpoint and coloring scheme are as in a. The observed oxidatively modified residues in spinach were mapped onto their corresponding locations on the LHC II associated with the *Zea mays* PS I-LHC I-LHC II supercomplex structure (PDB: 5ZJI, (Pan et al. 2018)) using Pymol (DeLano 2002). Oxidized amino acid residues are shown as spheres. Surface residues are shown in light brown and pink, respectively. Buried residues are shown in dark brown and red, respectively. In a, ^3^Thr of Lhcb2 is shown as spheres and colored by element. In b, Phospho-^3^Thr of Lhcb2 is shown as spheres and colored by element. Pigments are shown as sticks, chl in green and carotenoids in orange.

The majority of oxidatively modified residues in the LHC II isolated from both PS II membranes and PS I-LHC I-LHC II membranes, however, are buried (62% and 71%, respectively) in the hydrophobic environment at the interior of the Lhcb1 and Lhcb2 proteins and are not exposed to the bulk solvent. Since most ROS are hydrophilic (O2^-•^., H_2_O_2_, and .^•^OH), we would not expect these to diffuse from the bulk solvent to the hydrophobic interior of these proteins. We hypothesize that a subset of these observed modified residues would be in close proximity to the site(s) of ROS production within LHC II. Most of the buried oxidized residues in the hydrophobic cores of these subunits lie along the helix A and B axis (Fig. 4a, b). In these locations the modified residues are in very close proximity to the light-harvesting chls. For instance, in the Lhcb1 isolated from PS I-LHC I-LHC II membranes (Fig 4b), the observed oxidized buried residues are, on average, located 5.3 ± 1.7 Å (n=16) from the closest chl molecules.

Since ^1^O_2_ can be produced by the interaction of ^3^chl* with ^3^O_2_ by intersystem crossing, protective mechanisms are in place to prevent this from occurring frequently. The carotenoids present in LHC II can both physically quench ^3^chl* and ^1^O_2_ (Triantaphylidès and Havaux 2009) and also interact chemically with ^1^O_2_, forming carotenoid hydroperoxides and other oxidation products (Havaux 2013). In both instances, the ^1^O_2_ is prevented from damaging the protein matrix. Additionally, Non-Photochemical Quenching, NPQ, (Johnson and Ruban 2009) and LHC II aggregation (Natali et al. 2016) would mitigate the formation of ^1^O_2_ by shortening the lifetime of ^1^chl* (first singlet excited state of chl), thus minimizing the probability of ^3^chl* formation. However, these processes are not expected to be 100% efficient. If ^1^O_2_ is formed it may either exit the LHC II (possibly damaging other cellular components) or may directly oxidize the Lhcb proteins.

*In vitro*, LHC II trimers and monomers are known to produce ^1^O_2_ which causes significant protein damage (including cleavage) in detergent micelles (Zolla and Rinalducci 2002; Rinalducci et al. 2004), with ^1^O_2_ production being exacerbated when the LHC II is incorporated into lipid bilayers (Lingvay et al. 2020). Our results indicate that *in vivo* LHC II trimers also appear to produce ROS, most likely ^1^O_2_. Given that Lhcb proteins exhibit long ½ lives *in vivo* (∼10 hrs for *Lemna* and >24 hrs for *Phaseolus* (Slovin and Tobin 1982; Anastassiou and Argyroudi-Akoyunoglou 1995), it would be expected that each LHC II trimer may produce numerous ^1^O_2_ molecules during its ^2^lifetime. This could provide an ample opportunity for the oxidative modification of the Lhcb proteins. It should be noted that proteolysis-induced Lhcb turnover is exacerbated under high-intensity light conditions (Lindahl et al. 1995; Yang et al. 1998; Yang et al. 2000). This could be a response to increased oxidative damage to the Lhcb proteins.

While, overall, the oxidative damage observed for the LHC IIs associated with PS II membranes and PS I-LHC I-LHC II membranes appear similar, there are some important differences. First, as noted above, more surface-exposed residues are modified from the LHC II isolated from PS II membranes. Additionally, more buried oxidative modifications are found in the LHC II isolated from the PS I-LHC I-LHC II membranes (Fig. 4a,b). What could account for these differences, since these appear to indicate that the two LHC II preparations represent different populations *in vivo*ã PS II membranes represent the population of PS II which is found in the core regions of grana stacks and these lack grana margins. These are very similar regardless of isolation by either detergent treatment (Berthold et al. 1981; Dunahay et al. 1984) or mechanical isolation (Veerman et al. 2007; Danielsson and Albertsson 2009). The PS II located in this region produce a large amount of oxygen and, since PS II is the principal source of ROS in the chloroplast, would be expected to be a primary site of ROS-induced oxidative damage. Since many of the sites suggested for ROS production by PS II (Pheo_D1_, Q_A_, non-heme iron, and, possibly Q_B_) are proximal to the stromal-side of the membrane, release of ROS from these sites may preferentially damage the stromal-exposed regions on LHC II. While the Mn_4_O_5_Ca cluster, located on the lumenal-side of the membrane, may also produce ROS (principally .OH and H_2_O_2_ via the partial oxidation of water), .OH is extremely short lived and would not be expected to diffuse the distance required to modify LHC II, while H_2_O_2_ has relatively low reactivity. The stromal-exposed regions on the LHC II associated with PS I-LHC I-LHC II membranes would be less susceptible to ROS modification since PS I is a relatively minor source of ROS, probably producing significant amounts of these species only under certain environmental conditions (fluctuating light, high light intensities at low temperatures, acceptor-side limitations, etc.).

The observation that the LHC II associated with PS I-LHC I-LHC II membranes exhibit more buried oxidized residues may be explained by two factors. First, the LHC II associated with the PS I-LHC I supercomplex transfers energy relatively slowly (50-70 ps) to the reaction center (Bos et al. 2017). This slow excitation energy transfer rate increases the lifetime of ^1^chl* and increases the probability that ^3^chl* will form by intersystem crossing followed by ^1^O_2_ formation. A similar explanation may explain the high frequency of buried oxidized residues identified in LHC I (Kale et al. 2020). LHC I also transfers excitation energy slowly to the reaction center (∼45 ps) due to the presence of “red” chls in Lhca3 and Lhca4 (for a review, see (Croce and van Amerongen 2013)) as these serve as low energy sinks. While LHC II does not contain “red” chls, the slow excitation energy transfer rate from LHC II to the core antenna of PS I may be due to the absence of short chl-chl distances between LHC II and the core antennae chls which are principally associated with PsaA, PsaB, PsaK and PsaO, on the LHC II side of the PSI-LHC I supercomplex, as suggested for *Chlamydomonas* (Le Quiniou et al. 2015). The closest edge-to-edge chl-chl distances in *Z. mays* between Lhcb2 and PsaO, and between Lhcb1 and PsaK, is ∼8.0 Å (center-center chl-chl distances, 17.9 Å and 18.7 Å, respectively) (Pan et al. 2018).

Additionally, while 1-3 LHC II trimers (species-dependent) appear to be constituently associated with PS I (Chukhutsina et al. 2020; Croce 2020), additional LHC II trimers (up to 5, total (Bell et al. 2015)) may associate with the photosystem in State 2. During transit from PS II to PS I, these trimers are not associated with a reaction center and, consequently, the absorption of photons will lead to the formation of ^1^chl* which cannot be quenched photochemically. As noted above, subsequent formation of ^3^chl* would be expected to occur occasionally even in the presence of the various quenching mechanisms (carotenoids, NPQ, and LHC II aggregation). This would increase the probability of ^1^O_2_ formation with the concomitant possibility of protein oxidative damage.

The LHC II trimers which predominantly participate in state transitions are L-trimers, as it has been shown that the S- and M-trimers do not, in large measure, dissociate from the C2S2M2 complex upon phosphorylation (Wientjes et al. 2013a; Kyle et al. 1983). Consequently, in our study the LHC II which are associated with PS II membranes and isolated from the core region of grana stacks, are expected to consist primarily of S- and M-trimers. The LHC II associated with PS I-LHC I-LHC II membranes, which are isolated from the grana margins and/or the stroma lamella (Chukhutsina et al. 2020), consist primarily of the mobile L-trimers and trimers constitutively associated with PS I. Our current study indicates that these two populations of LHC II trimers exhibit differential oxidative modification patterns. The LHC II associated with the C2S2M2 supercomplex is energetically coupled with the PS II reaction center a very high proportion of the time, which minimizes the formation of ^3^chl* and lowers the probability of ^1^O_2_ formation and subsequent oxidative damage to buried residues. Additionally, because of their close proximity and stable association with PS II, which produces a variety of hydrophilic ROS, the surface-exposed domains would be expected to exhibit enhanced rates of oxidative modifications. The L-trimers, during state transitions, are uncoupled from reaction centers, increasing the probability of ^3^chl* formation and increasing the probability of oxidative damage to buried residues by ^1^O_2_. The trimer(s) constitutively associated with the PS I-LHC-I supercomplex appears to exhibit slow energy transfer rates to the reaction center (Chukhutsina et al. 2020; Bos et al. 2017). This may also lead to enhanced ^3^chl* formation. Our study also suggests that L-trimers must exchange infrequently with S- or M-trimers. If such exchange occurred at significant rates, and given the long ½ life of Lhcb components of LHC II, one would expect that the oxidative modification patterns would be quite similar, if not identical, between the LHC II isolated from these two different sources.

Finally, it should be noted that the pattern of oxidative modification observed for the Lhcb proteins in LHC II are remarkably similar to the oxidative modification pattern observed for the Lhca proteins associated with the PS I-LHC I supercomplex (Kale et al. 2020). This is illustrated in Fig. 5. As noted above, the buried oxidative modifications observed for Lhcb1 protein associated with PS I-LHC I-LHC II are primarily associated with the A and B helixes (Fig. 5a). In the Lhca3 associated with the LHC I of the PS I-LHC I supercomplex (Fig. 5b) a very similar pattern is observed, with most of the buried oxidative modifications associated with the A and B helices of this component. This is also true of the other Lhca proteins associated with LHC I. These oxidative modifications are also in close proximity to the light-harvesting chls (4.3 ± 1.1 Å, n=13). There is no statistical difference between these values and those observed for the distance between oxidized residues in Lhcb1 and chl (5.3 ± 1.7 Å, n=16). The similarity in the locations of the oxidative modifications in Lhcb1 and Lhca3 and their close proximity to light-harvesting chls suggests a common mechanism for the production of this observed damage, i.e., ^1^O_2_ production by unquenched ^3^chl*. Earlier, it was determined that the LHC I associated with the PS I-LHC I supercomplex is rapidly lost during PS I photoinhibition (Alboresi et al. 2009; Hui et al. 2000). This appears analogous to the loss of Lhcbs (Lindahl et al. 1995; Yang et al. 1998; Yang et al. 2000) associated with PS II under high intensity light illumination. In both instances, increased oxidative damage may trigger enhanced proteolytic degradation of these proteins.

**Figure 5.**
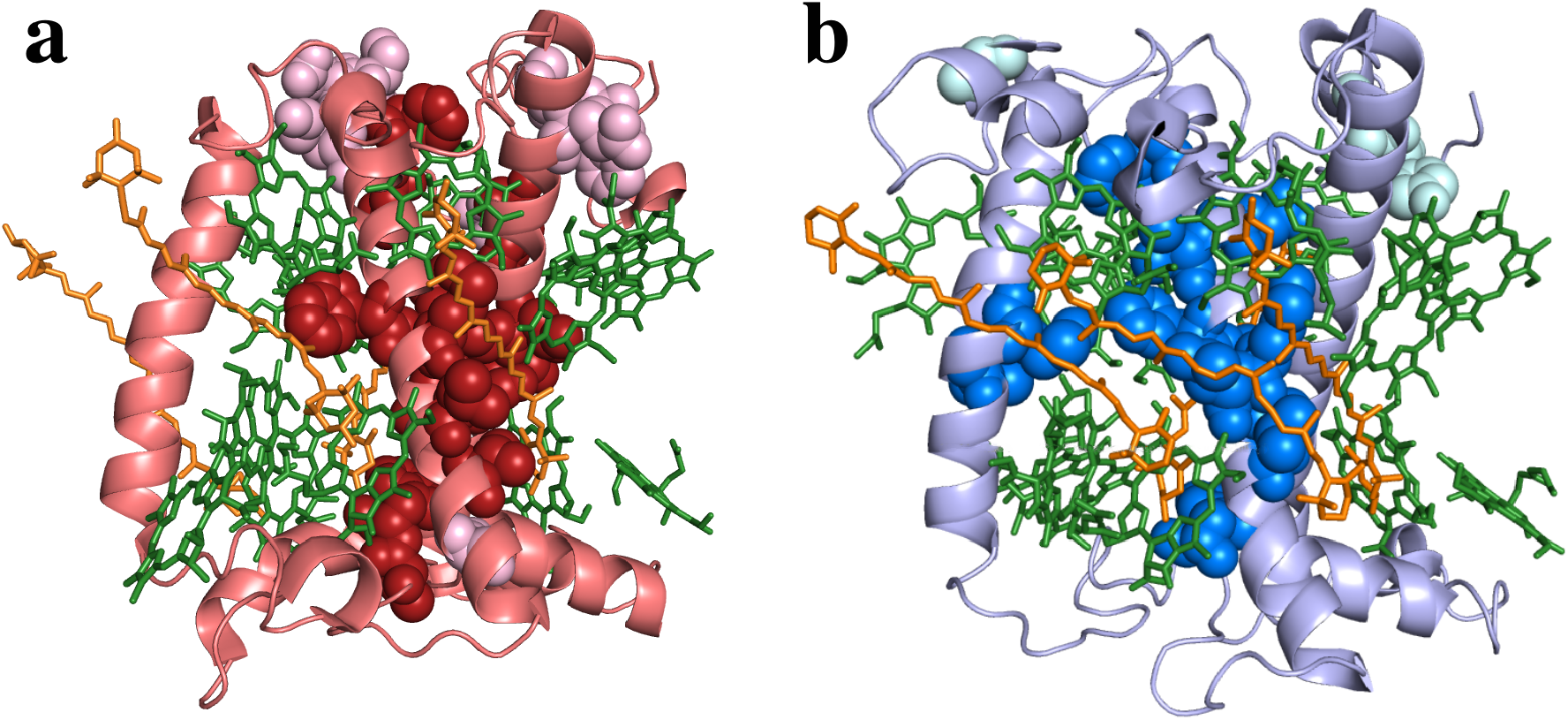
Comparison of the Oxidative Modifications found in Lhcb1 Associated with PS I-LHC I-LHC II Membranes and Lhca3 Associated with the PS I-LHC I Supercomplex. a, Lhcb1 associated with PS I-LHC I-LHC II Membranes. The observed oxidatively modified residues in spinach were mapped onto their corresponding locations on the LHC II associated with the *Zea mays* PS I-LHC I-LHC II supercomplex structure (PDB: 5ZJI, (Pan et al. 2018)). b, Lhca3 associated with the PS I-LHC I supercomplex. The oxidatively modified residues in spinach (Kale et al. 2020) were mapped onto their corresponding locations on Lhca3 associated with the *Pisum sativum* PS I-LHC supercomplex structure (PDB: 5L8R, (Mazor et al. 2017)). Viewpoint for both panels is within the plane of the membrane. Proteins are shown as cartoon representations. Oxidized amino acid residues are shown as spheres. Surface residues are shown in light shades while buried residues are shown as darker shades. Pigments are shown as sticks, chl in green and carotenoids in orange. These visualizations were produced using Pymol (DeLano 2002)

## Conclusions

In this communication we have documented that the LHC II associated with PS II membranes and PS I-LHC I-LHC II membranes exhibit significant oxidative modification. While a subset of these oxidative modifications is located on these proteins’ surfaces that are exposed to the bulk solvent and potentially a variety of hydrophilic ROS generated from different sources, the majority of these oxidized residues are buried within the hydrophobic membrane protein matrix. We hypothesize that these modifications are the result of oxidative damage by ^1^O_2_ which is produced through the interaction of ^3^O_2_ with ^3^chl*. While somewhat similar, the overall pattern of oxidative modifications for the LHC II isolated from these two sources differs significantly in that a higher proportion of surface-exposed oxidative modifications are observed for the LHC II associated with PS II membranes while a higher proportion of buried oxidative modifications are observed for the LHC II associated with PS I-LHC I-LHC II membranes. Finally, the oxidative modifications which we previously observed for the Lhca proteins associated with LHC I (Kale et al. 2020) are very similar to those that we observe for the Lhcb proteins associated with LHC II. This observation suggests that the source of the ROS associated with the oxidative modifications of these two protein types arise from similar sources, namely from ^1^O_2_ production from the interaction of ^3^O_2_ with ^3^chl*.

## Acknowledgements

This work was solely supported by the United States Department of Energy, Office of Basic Energy Sciences grant DE-FG02-09ER20310 to TMB and LKF.

## Footnotes

1 Considering only one Lhcb1 and one Lhcb2 within the trimer.

2 For a **very approximate estimate**, consider an LHC II trimer which is not associated with a reaction center (for instance if it were moving between PS II and PS I during State 1 to State 2 transitions). Since each chl absorbs about 10 photons/sec (at 1800 **μ**moles photons/m^2^/sec) (Blankenship 2002) and each LHC II trimer contains 42 chl, each LHC II trimer absorbs about 420 photons/sec. If an LHC II trimer has a ½ life of 10 hrs, it would absorb 1.5 × 10^7^ photons. Assuming the ^3^chl*-quenching mechanisms (carotenoids, NPQ and LHC II aggregation, etc.) were 99.99% effective at quenching ^3^chl* and that 10% of the unquenched ^3^chl* formed ^1^O_2_, then ∼150 ^1^O_2_ could be formed over the lifetime of the LHC II. LHC II trimers connected to reaction centers would be predicted to produce significantly less ^1^O_2_.

